# Facilitation of hERG channels by blockers: a mechanism predicted to reduce lethal cardiac arrhythmias

**DOI:** 10.1101/341875

**Authors:** Kazuharu Furutani, Kunichika Tsumoto, I-Shan Chen, Kenichiro Handa, Yuko Yamakawa, Jon T. Sack, Yoshihisa Kurachi

## Abstract

Fatal cardiac arrhythmias are caused by some, but not all, drugs that inhibit the cardiac rapid delayed-rectifier current (*I*_Kr_) by blocking hERG channels. Here, we propose a novel mechanism that could make certain hERG blockers less proarrhythmic. Several drugs that block hERG channels, yet have favorable cardiac safety profiles, also evoke another effect; they increase the current amplitude upon low-voltage depolarization (facilitation). Voltage-clamp recordings of hERG block and facilitation by nifekalant, a Class III antiarrhythmic agent, constrained a model of human cardiac *I*_Kr_. In human ventricular action potential simulations, nifekalant showed its therapeutic ability to suppress ectopic excitations, with or without facilitation. Without facilitation, excessive *I*_Kr_ block evoked early afterdepolarizations, which cause lethal arrhythmias. Facilitation prevented early afterdepolarizations at the same degree of block by increasing *I*_Kr_ during repolarization phase of action potentials. Thus, facilitation is proposed to reduce the arrhythmogenic risk of hERG blockers.

**Abbreviations:** AP: action potential; APD: action potential duration; APD_90_: action potential duration measured at 90% repolarization; EAD: early afterdepolarization; hERG: human ether-ago-go-related gene; *I*_CaL_: L-type Ca^2+^ channel current; *I*_net_: net ionic current; *I*_K1_: inward-rectifier potassium current; *I*_Kr_: rapid component of the delayed-rectifier potassium current; *I*_Na_: sodium current; ORd: O’Hara-Rudy dynamic

## Introduction

A rapid component of the delayed-rectifier potassium current (*I*_Kr_) plays an important role in the repolarization of the cardiac action potential (AP). *I*_Kr_ is especially important in ventricular muscle of the hearts of large mammals. Certain blockers of *I*_Kr_ are class III antiarrhythmic agents, used to treat ventricular tachyarrhythmias (*Sanguinetti and Jurkiewicz 1990, Vaughan Williams 1992, Zeng et al. 1995, Clancy et al. 2003*). Blockers of *I*_Kr_ prolong the AP duration (APD) and effective refractory period (ERP) to suppress premature ventricular contraction. Blockade of *I*_Kr_ also is a side-effect of many drugs. Blockers of *I*_Kr_ can cause acquired long QT syndrome and the life-threatening ventricular tachyarrhythmia called “*torsades de pointes*” (*Surawicz 1989, Sanguinetti et al. 1995, Roden 2000, Sanguinetti and Tristani-Firouzi 2006, Roden 2008*). Emphasis on detecting *I*_Kr_ blockade early in drug discovery (*ICH 2005*) has contributed to the successful removal of *torsades de pointes* risk for new chemical entities. However, there is concern that promising new drug candidates are being unnecessarily discarded because they benignly block *I*_Kr_ (*Sager et al. 2014, Gintant et al. 2016*). Indeed, numerous examples exist of drugs that block *I*_Kr_ and prolong the QT interval but have little proarrhythmic risk (*De Ponti et al. 2001, Redfern et al. 2003*).

The human *ether-a-go-go-related gene* (hERG) encodes the ion channel through which *I*_Kr_ current passes (*Sanguinetti et al. 1995*). We have found that some hERG blockers also increase hERG currents at potentials close to the threshold for channel activation. We refer to this electrical phenomenon as ‘facilitation’ (*Hosaka et al. 2007, Furutani et al. 2011, Yamakawa et al. 2012*). A series of hERG blockers with lower proarrhythmic risk were found to evoke facilitation of hERG (*Hosaka et al. 2007, Furutani et al. 2011, Yamakawa et al. 2012*). The correlation between clinical safety and facilitation lead to our current hypothesis: *facilitation by hERG blockers increases I_Kr_ during cardiac AP repolarization and thereby decreases their proarrhythmic risk.* Stringently testing this hypothesis requires a system where facilitation can be selectively removed from a hERG blocker’s mechanism. In the present study, we test the aforementioned facilitation hypothesis with a mathematical model of the actions of nifekalant, a hERG channel blocker that induces facilitation and has been used safely in the treatment of life-threatening ventricular tachyarrhythmias (*Nakaya et al. 1993, Takenaka et al. 2001, Igawa et al. 2002, Sato et al. 2017*), on the APs of human ventricular myocytes.

## Results

### AP voltage stimuli induce facilitation by nifekalant

The degree of facilitation of *I*_Kr_ depends on the dose of hERG blocker and the membrane’s history of voltage changes. Facilitation by blockers, with the exception of amiodarone derivative KB130015 (*Gessner et al. 2010*), requires a preceding strong depolarization as a conditioning stimulus (*Carmeliet 1993, Jiang et al. 1999, Hosaka et al. 2007, Furutani et al. 2011, Yamakawa et al. 2012*). To test whether cardiac APs are sufficient to stimulate facilitation, we applied an AP clamp protocol to hERG channels. Repeated stimulation with a human ventricular myocyte waveform at 1 Hz induced facilitation of nifekalant-treated hERG channels (*Figure 1*). The facilitation effect on hERG current saturated as the number of APs increased, indicating facilitation reached 90% of its steady state level within 20 heartbeats (*Figure 1C*). The induction of facilitation by cardiac AP stimuli suggests that facilitation is potentially physiologically relevant.

**Figure 1.**
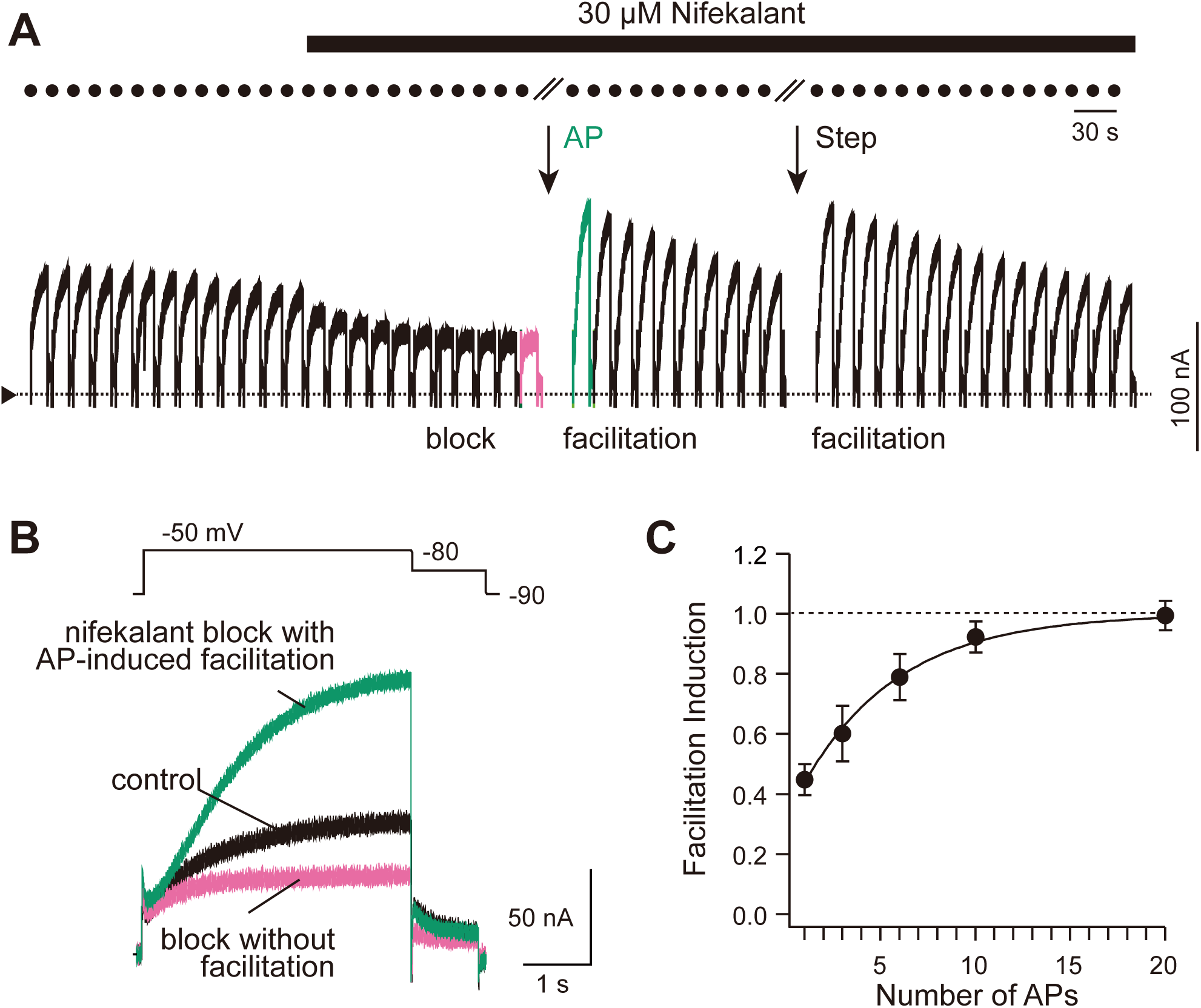
Cardiac APs induce hERG facilitation with nifekalant. (**A, B**) Representative cell currents from hERG channels in *Xenopus* oocytes evoked by a test pulse from holding potential of −90 mV to −50 mV before and after AP stimulation (1 Hz, 20×, AP waveform is the same as *Figure 2H*) in the presence of 30 μM nifekalant. (**C**) AP-facilitation relation. The fraction of facilitation induced by repeating APs are normalized to the fraction induced by the +60 mV conditioning pulse. Experimental data are means ± SEM (*n* = 8-15). The curve fit indicates exponential increase in the facilitation fraction by AP stimulation (Facilitation = 1.02 − 0.70·exp(−(#APs)/*τ*), *τ* = 5.45 ± 0.02).

### Facilitation can be incorporated into cardiac electrophysiology modeling

To understand how facilitation could impact cellular electrophysiology mechanisms specific to human ventricular myocytes, we used the O’Hara-Rudy dynamic (ORd) human ventricular AP model (*O’Hara et al. 2011*). We modified the *I*_Kr_ formula in the ORd model to reproduce hERG channel block by nifekalant, with or without facilitation. This modified *I*_Kr_ formula was constrained by currents recorded from HEK cells expressing hERG (*Figure 2A-C*; *Supplemental Figure 1*). *Figure 2* shows the agreement of our *I*_Kr_ model with hERG current from HEK cells and its modulation by nifekalant. Left panels of *Figure 2D, E* show experimental hERG currents blocked with 100 nM nifekalant, before (magenta traces) and after the induction of facilitation effect by a conditioning pulse (green traces). After the induction of the facilitation effect, nifekalant enhanced the current induced by a −30 mV pulse (*Figure 2D*), but it still suppressed the current induced by a +30 mV pulse (*Figure 2E*). The current enhancement is due to a modification of channel activation gating that shifts the voltage dependence to more hyperpolarized membrane potentials (*Hosaka et al. 2007, Furutani et al. 2011*). The voltage dependence of the hERG current in the facilitation condition could be described as the sum of two Boltzmann functions reflecting two populations of hERG currents having different activation voltage dependences (*Furutani et al. 2011*). The high-*V*_1/2_ fraction of the channel population has biophysical characteristics typical for the hERG channel, while the low-*V*_1/2_ fraction is ‘facilitated’. The facilitated *V*_1/2_ was shifted negative by −26.5 mV (*Table 1*). This shift was consistent with that of hERG channels expressed in *Xenopus* oocytes (*Furutani et al. 2011*). Thus, the facilitation effect could be explained by a fraction of channels opening with a negatively shifted activation curve (*Figure 2F, G*). Inclusion of the facilitation effect in the *I*_Kr_ model reproduced measurements of hERG currents from voltage-step experiments (*Figure 2D-G*), and over a range of concentrations (*Supplemental Figure 2*). In AP clamp experiments and simulations, our *I*_Kr_ model also predicted the effects of nifekalant on hERG currents during AP (*Figure 2H*), indicating that the model describes the block and facilitation effects of nifekalant sufficiently to allow us to assess the impact of facilitation in simulations of human ventricular myocyte electrophysiology.

**Table 1.** Effects of nifekalant on hERG channels

| Dose-response |  |  |
| --- | --- | --- |
|  | IC<sub>50</sub>/EC<sub>50</sub> (nM) | hill coefficient |
| Block | 144.92 ± 16.00 | 1.15 |
| Facilitation | 92.84 ± 7.71 | 1.50 |
| Shift of the hERG activation curve by facilitation effect |  |  |
|  | ΔV<sub>1/2</sub> (mV) |  |
| HEK293 stably-expressing hERG | -26.5 |  |
| HEK293 transiently-expressing hERG | -24.9 |  |
| HEK293 transiently-expressing hERG with KCNE2 | -24.4 |  |
| X. Oocyte transiently-expressing hERG | -26.6 (2) |  |

**Figure 2.**
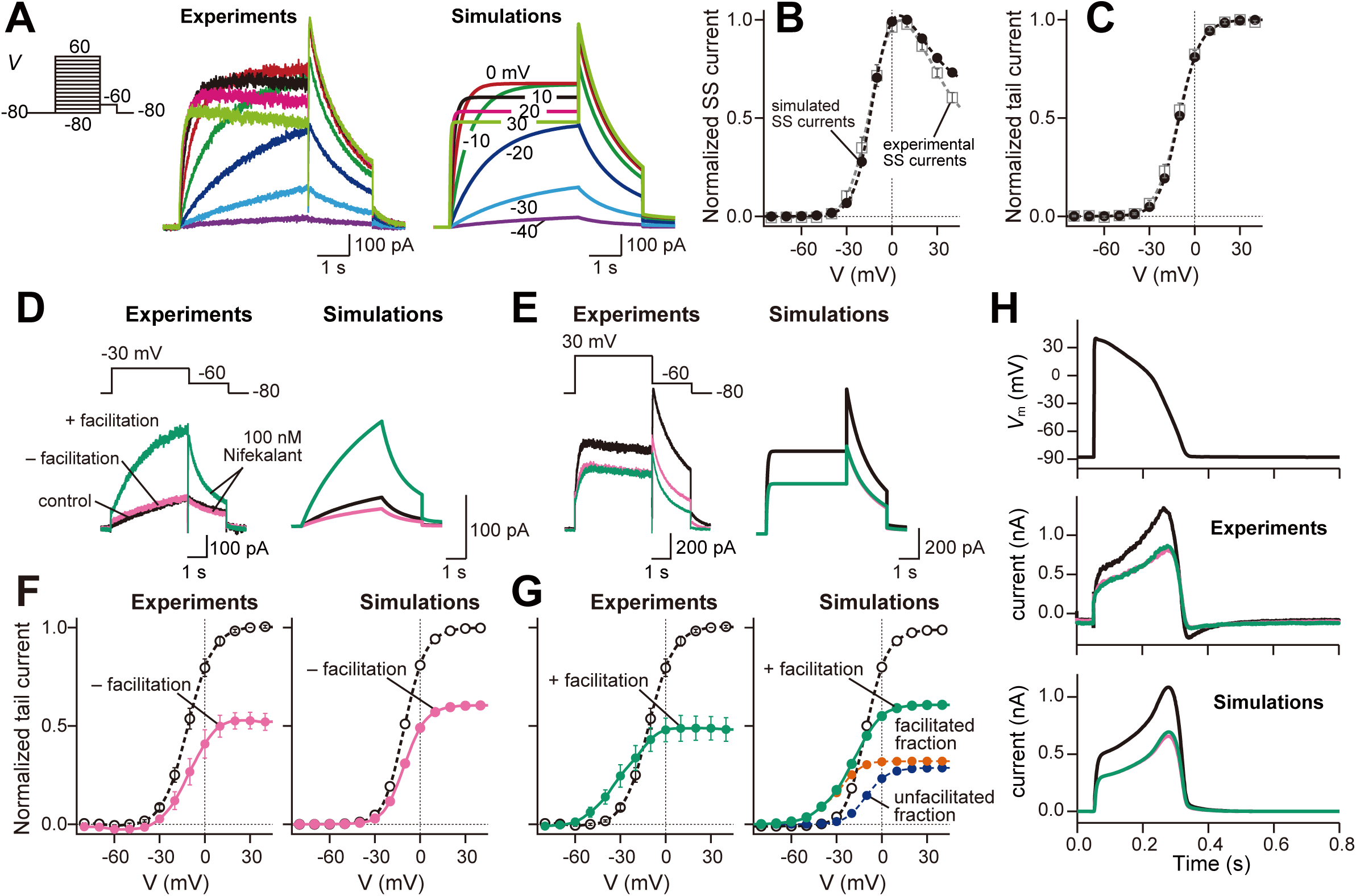
Experimental and simulated macroscopic hERG/*I*_Kr_ currents as modified by nifekalant. (**A-C**) The macroscopic hERG/*I*_Kr_ currents in response to voltage-clamp pulses from −80 mV to +60 mV in 10 mV increments from a holding potential of −80 mV; representative traces (**A**), the relationship between membrane voltage and the steady-state current amplitude and membrane voltage (**B**) and the tail current amplitude (activation curve) (**C**). Experimental data are means ± SEM (*n* = 11). (**D-G**) Simulated effects of nifekalant on hERG/*I*_Kr_ currents. The model assumes two populations of channels, with or without facilitation effect by nifekalant (100 nM). The *V*_1/2_ of activation for the facilitated fraction of channel was ~ −31 mV, almost 26 mV negative to that of control channel (see also Materials and Methods). (**D**, **E**) The macroscopic hERG/*I*_Kr_ currents in response to voltage-clamp pulses from −80 mV to −30 mV (**D**) or +30 mV (**E**). (**F**, **G**) The relationship between the tail current amplitude and membrane voltage before (**F**) and after (**G**) the induction of facilitation effect. Experimental data are means ± SEM (*n* = 5). (**H**) The macroscopic hERG/*I*_Kr_ currents in response to cardiac AP. In **D-H**, black, red, and green solid lines indicate with control, block with facilitation, and conventional block (block without facilitation), respectively. Under the condition in the block with facilitation, *I*_Kr_ comprises two fractions of *I*_Kr_ (see also main text), i.e., facilitated and unfacilitated fractions of *I*_Kr_. In **G** right panel, orange and cyan dashed lines represent the facilitated and unfacilitated fractions of *I*_Kr_, respectively. Experimental data are means ± SEM (*n* = 6-11).

### Facilitation selectively increases I_Kr_ during late phase-2 and phase-3 repolarization of cardiac APs

To understand the role of facilitation in the cardiac AP, we investigated the influence of facilitation on AP morphology. When endocardial APs are evoked with 1 Hz pacing, 100 nM nifekalant prolonged the APD measured at 90% repolarization, APD_90_, (334.4 ms, vs. 257.4 ms in the control condition, see *Figure 3A*), in agreement with experimental results in human ventricular myocytes (*Jost et al. 2005*). The AP was slightly further prolonged in simulations with *I*_Kr_ block without facilitation (342.0 ms APD_90_, compare green and magenta traces in *Figure 3A*). In the model, 100 nM nifekalant blocks 40% of total *I*_Kr_ at voltages > 0 mV with or without facilitation (*Figure 2G*). When facilitation is implemented in the model, 32% of the hERG channels enter a facilitated state such that they produce substantially more *I*_Kr_ than control conditions when the potential is between −20 and −50 mV (*Figure 2G*). There was little difference between the voltage-dependent activation variables in the *I*_Kr_ model during phase-1 and phase-2 of the AP (*x*_r1_ and *x*_r2_ for the unfacilitated channels and facilitated channels, respectively), indicating *I*_Kr_ behaved similarly (*Figure 3B, C*). During late phase-2 and phase-3 repolarization, an increase can be seen in *I*_Kr_ (asterisks in *Figure 3A-C*). However, in block conditions with or without facilitation, the APs were terminated by increase in the inward-rectifier potassium current (*I*_K1_)-mediated repolarization current (*Figure 3D*) after a short time, before much prolongation of the AP by facilitation. Thus, in this model of a healthy heart under unstressed conditions, facilitation by this hERG blocker selectively increases *I*_Kr_ during late phase-2 and phase-3 repolarization, but this has only a subtle effect on the cardiac AP.

**Figure 3.**
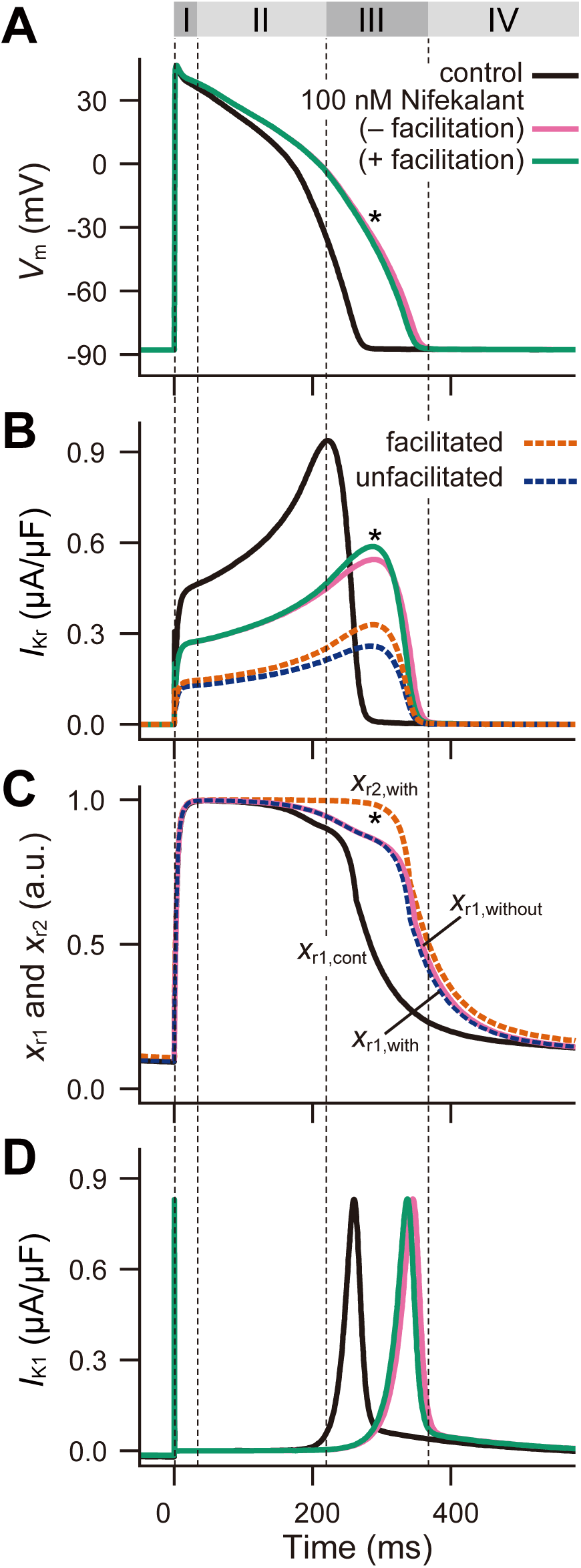
Effect of *I*_Kr_ facilitation on cardiac AP. (**A**) Simulated APs in an endocardial ventricular myocyte, (**B**) *I*_Kr_, (**C**) activation state values for unfacilitated (*x*_r1_) and facilitated (*x*_r2_) fractions of *I*_Kr_, and (**D**) *I*_K1_ during APs with 100 nM nifekalant. Through **A** to **D**, black, green, and magenta solid lines indicate with control, block with facilitation, and conventional block (block without facilitation), respectively. Roman numerals above **A** indicate the phases of the AP in the case of with facilitation. a.u. indicates arbitrary unit.

### Facilitation relieves reverse frequency-dependence of APD prolongation

Class III arrhythmic agents are hERG blockers that are used clinically to suppress ventricular tachyarrhythmias (*Sanguinetti and Jurkiewicz 1990, Vaughan Williams 1992, Sanguinetti and Tristani-Firouzi 2006*). To suppress tachyarrhythmias without provoking *torsades de pointes*, *I*_Kr_ block would ideally be use-dependent and prolong APD only in response to high frequency stimulation (*Surawicz 1989, Hondeghem and Snyders 1990*). Previous studies reported that actions of sotalol (*Hafner et al. 1988, Hondeghem and Snyders 1990*) and dofetilide (*Tande et al. 1990*), trend against this antiarrhythmic ideal by prolonging APDs more at low than high frequencies, showing reverse frequency-dependence. Notably, sotalol and dofetilide are hERG blockers that do not induce facilitation (*Furutani et al. 2011, Yamakawa et al. 2012*). Nifekalant also showed reverse frequency-dependence, but it was not as marked (*Nakaya et al. 1993, Cheng et al. 1996, Igawa et al. 2002*). To determine why, we used our model to test the frequency-dependent effects of nifekalant on APD. The left panel of *Supplemental Figure 3* shows changes in the APD_90_ with 100 nM nifekalant at stimulation frequencies from 0.2 to 2 Hz. Without facilitation (magenta trace in *Supplemental Figure 3*, left), the prolongation of APD was more effective at lower stimulation frequencies, resulting in a reverse frequency-dependence. When facilitation was included in the model, the reverse frequency-dependence became slightly weaker at low frequencies (green trace in *Supplemental Figure 3*, left). APD_90_ with facilitation was 5.5 ms shorter than without at 2.0 Hz, 7.7 ms shorter at 1.0 Hz, and 10.7 ms shorter at 0.2 Hz (*Supplemental Figure 3*, right). This suggests facilitation can partially relieve the reverse frequency-dependence that is a risk factor for *torsades de pointes*.

### I_Kr_ facilitation has greater impacts on repolarization in a failing heart model

Heart failure patients are at risk for malignant ventricular arrhythmias. Clinical and theoretical data has shown that the APs in heart failure patients at slow and modest heart rates are destabilized compared with healthy subjects (*Bayer et al. 2010*). A more recent study indicated the APs of the heart failure models exhibit stronger rate-dependence when compared with the APs of the normal model, and can generate early afterdepolarizations (EADs) and alternans at modest pacing rates (*Elshrif et al. 2015*). These features are thought to trigger ventricular arrhythmias (*Roden 2000, Thomsen et al. 2003, Nattel et al. 2007, Roden 2008, Weiss et al. 2010*). To determine whether facilitation might have a more critical impact on failing hearts, we investigated the frequency-dependent effects of *I*_Kr_ facilitation on the simulated AP in a heart failure model (*Figure 4*) (*Elshrif et al. 2015*). APs in the heart failure model were prolonged from the control (non-heart failure) model (compare black traces in *Figure 4B*, *Supplemental Figure 3*), and were further prolonged by *I*_Kr_ blockade. The effect of facilitation was more prominent with low frequency pacing of the heart failure model, and the reverse frequency-dependence of APD prolongation was more dramatically attenuated (compare *Figure 4A*, *Supplemental Figure 3*, left) because the delayed repolarization at low frequency pacing prolongs the latency before facilitation increases the *I*_Kr_ current. APD_90_ during block with facilitation was 16.5 ms shorter than without facilitation at 2.0 Hz, 36.3 ms shorter at 1.0 Hz, and 53.0 ms shorter at 0.2 Hz (*Figure 4B*), suggesting that the electrical remodeling in failing myocytes makes cellular responses more sensitive to *I*_Kr_ modulation.

**Figure 4.**
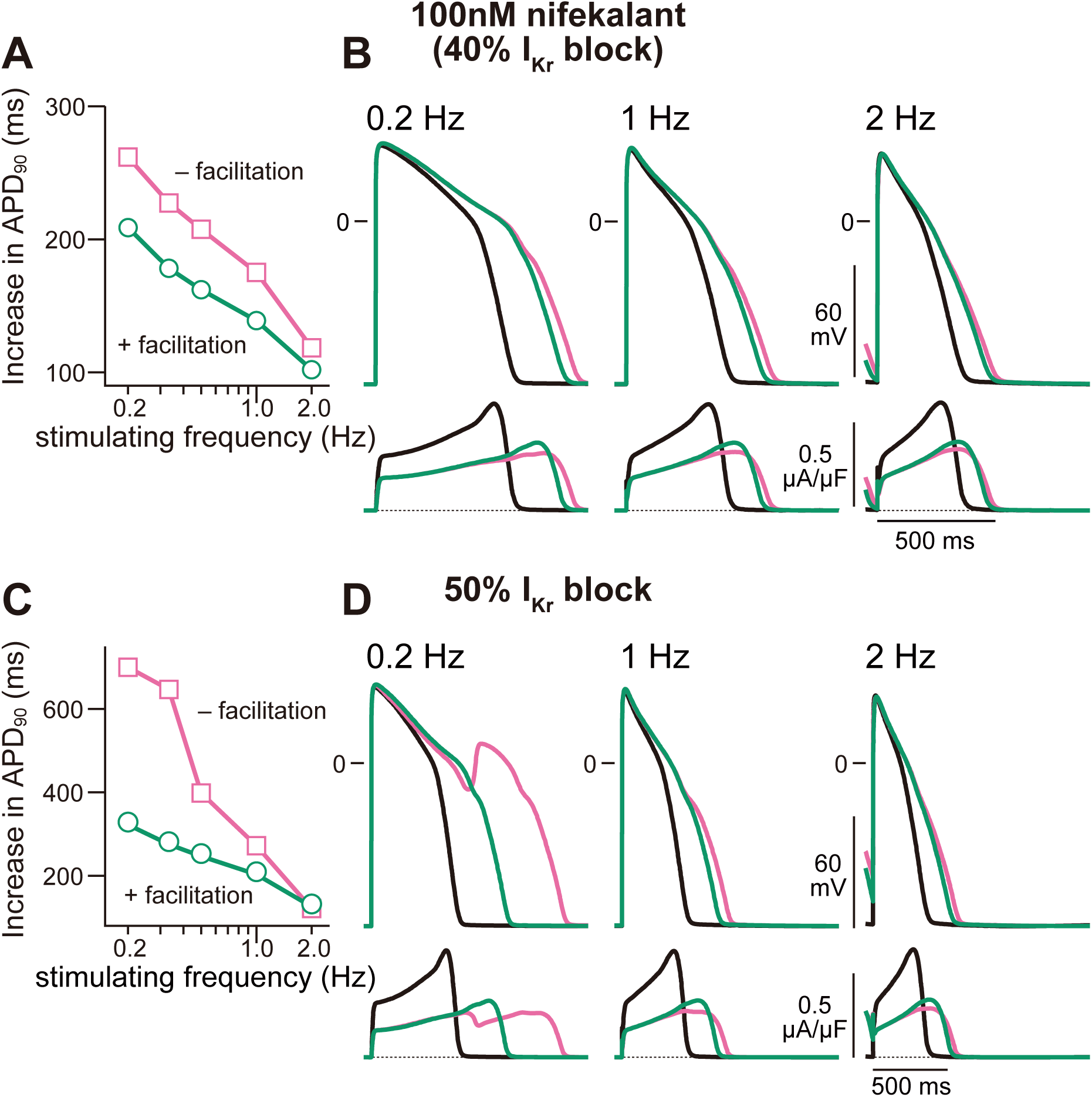
Frequency-dependent effect of nifekalant on APD. Increases in simulated APD_90_ from the control condition in the heart failure model with 100 nM nifekalant (40 % *I*_Kr_ block) (**A**) and 50% *I*_Kr_ block (**C**) at each stimulation frequency. Effects of *I*_Kr_ block and facilitation on the APs in the heart failure model at various simulation frequency with 100 nM nifekalant (40 % *I*_Kr_ block) (**B**) and 50% *I*_Kr_ block (**D**).

### I_Kr_ facilitation prevents EAD

Excessive AP prolongation by a hERG blocker showing strong reverse frequency-dependence creates an electrophysiological environment that favors the development of EAD at low stimulating frequencies. Increasing *I*_Kr_ block from 40% to 50% caused further APD prolongation (*Figure 4C, D*). Without facilitation, the APD was dramatically prolonged with low frequency pacing (magenta trace in *Figure 4C*). We found that in the heart failure model paced at 0.2 Hz, EAD appeared at 50% *I*_Kr_ block without facilitation (magenta traces in *Figure 4D*), and facilitation suppressed these EADs (green traces in *Figure 4D*). Although this pacing is unphysiologically slow, these findings suggest hERG facilitation could have a more significant impact on arrhythmogenesis in a failing heart.

To determine the range of conditions over which facilitation could potentially suppress proarrythmic effects of hERG blockade, we extended analysis in the heart failure model. *Figure 5A-C* shows examples of steady-state AP trains in several *I*_Kr_ block conditions. *Figure 5D* shows the changes in APD_90_ as a function of the degree of *I*_Kr_ block. Below 40% block, the drug prolonged APD_90_ similarly with or without facilitation (*Figure 5A, D*). Without facilitation, APs were destabilized when *I*_Kr_ inhibition was > 48% (section sign in *Figure 5D*; APD_90_ = 890.0 ms), resulting in an alternating EADs and periodic EADs. With facilitation, EADs did not appear until 57% inhibition of *I*_Kr_ (pipes in *Figure 5D*; APD_90_ = 896.4 ms), indicating that *I*_Kr_ facilitation stabilized the AP. *Figure 5B, C* show examples of steady-state AP trains with 50% and 55% block of *I*_Kr_ (conditions indicated by dagger and double-dagger in *Figure 5D*, bottom), respectively. These results suggest the facilitation of *I*_Kr_ by blockers reduces proarrhythmic side effects by preventing the development of EADs. In the non-failing heart model, a similar antiarrhythmic effect of facilitation was observed with more extreme *I*_Kr_ blockade (*Supplemental Figure 4*).

**Figure 5.**
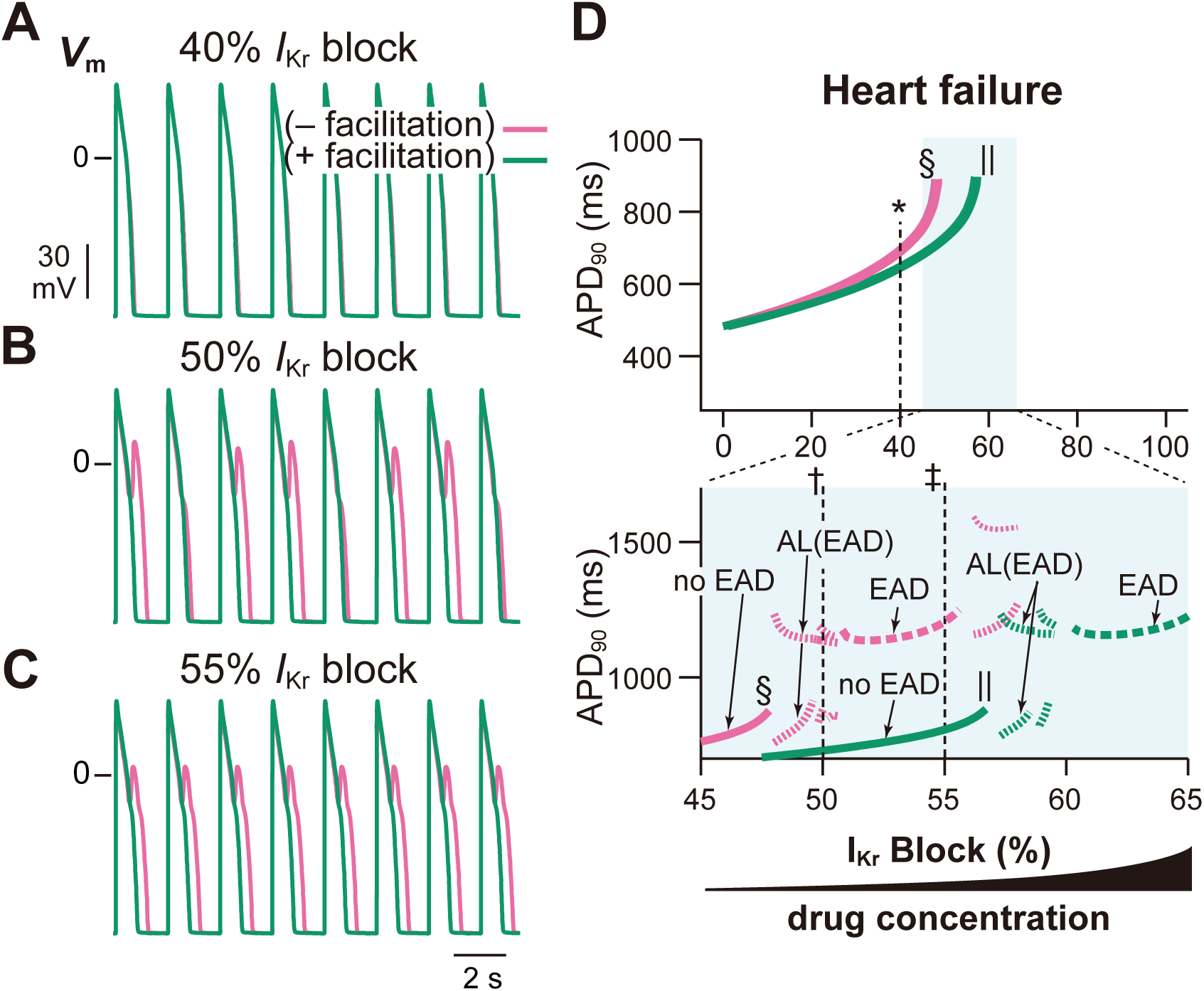
The effect of *I*_Kr_ facilitation on proarrhythmic risk. (**A-C**) The steady-state AP trains with 40% (**A**), 50% (**B**), and 55% *I*_Kr_ block (**C**) in heart failing model with and without facilitation. (**D**) Effect of *I*_Kr_ block and facilitation on the APD and the development of EADs in heart failure model. Asterisk, dagger, and double-dagger indicate the conditions of in **A-C**, respectively. Sections and pipes indicate the upper limits of *I*_Kr_ block where APs ware normally terminated. In the bottom panel of **B**, AL(EAD), alternated EAD (thin dashed trace); EAD, periodic EAD (bold dashed trace), see also main text.

*I*_Kr_ facilitation suppresses reactivation of L-type Ca^2+^ channels. In the heart failure model at 55% inhibition of *I*_Kr_ without facilitation (magenta lines in *Figure 6,* left), the repolarization delay caused by *I*_Kr_ reduction augmented the window current in L-type Ca^2+^ channel current (*I*_CaL_) during late AP phase-2 with each stimulation (*Figure 6C*, left). As a result, the evoked APs were gradually prolonged with each stimulation, leading to further augmentation of the *I*_CaL_ window current (*Figure 6C*, left) and AP prolongation (*Figure 6B*, left). Then, the stagnated repolarization caused more reactivation of L-type Ca^2+^ channels (arrowhead in *Figure 6C*, left), enhancing an inward current component of the net ionic current (*I*_net_) during late AP phase-2 (*Figure 6D*, left). This caused the inward-outward balance of *I*_net_ (asterisks in *Figure 6D*, left) that temporally interrupted AP repolarization, followed by a second rising phase of the AP that led to EAD. This *I*_Kr_ block-induced dysfunction eventually converged to a periodic EAD response (*Figure 5C* and *Figure 6A*, right column). In contrast, an increase in *I*_Kr_ due to facilitation (*Figure 6E*, right) enhanced an outward current component in *I*_net_ during late AP phase-2 and phase-3 (*Figure 6D*, right). This accelerated the membrane repolarization and completed the AP repolarization (*Figure 6B*, right). Thus, facilitation suppresses EADs by selectively amplifying *I*_Kr_ during late phase-2 and phase-3 of APs when the repolarization is in danger of stagnating. The facilitated *I*_Kr_ overwhelms the reactivating *I*_CaL_, preventing EAD.

**Figure 6.**
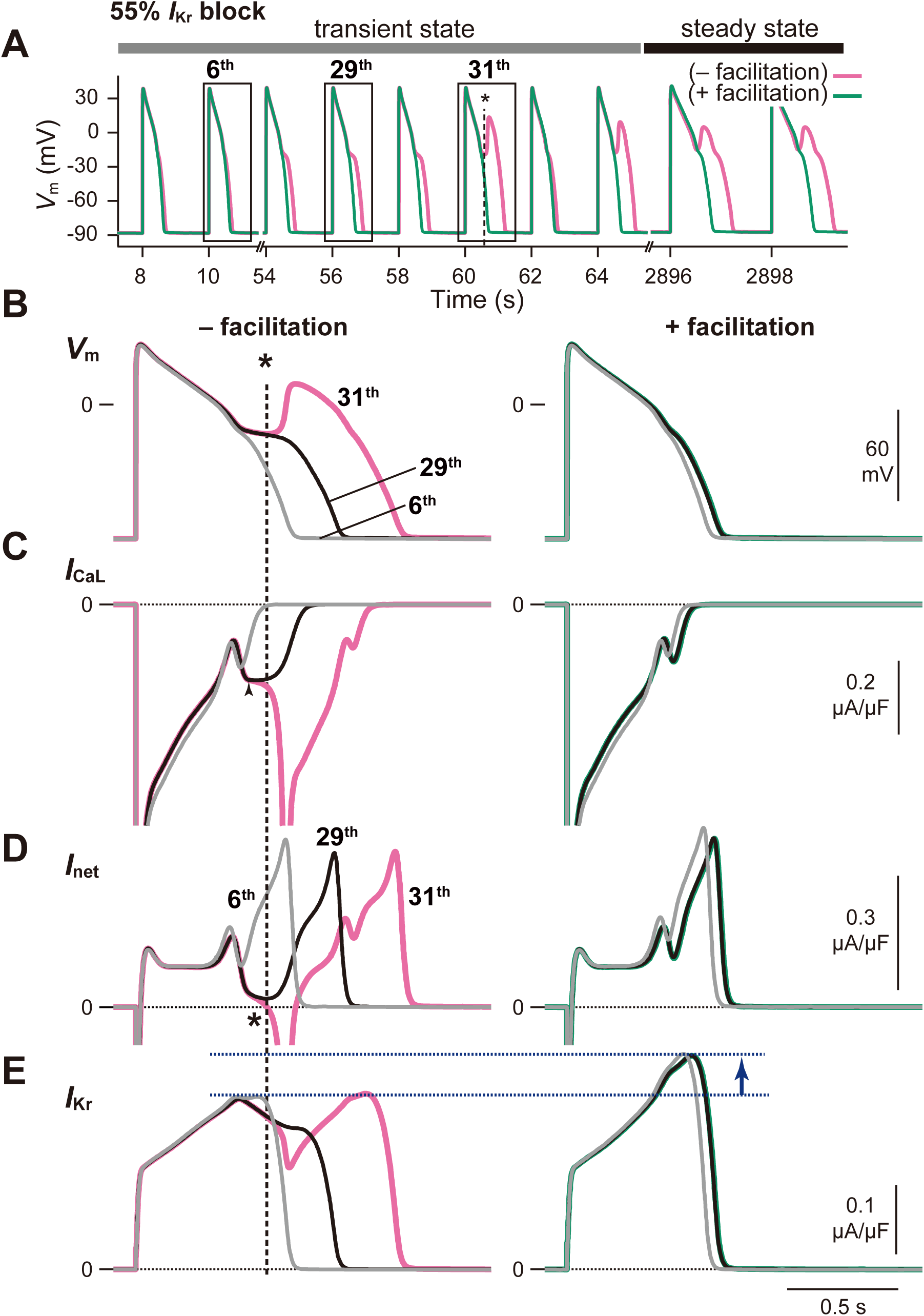
Ionic mechanism of EAD development by *I*_Kr_ block and the influence of *I*_Kr_ facilitation. Simulated APs and the changes in the membrane potential (*V*_m_) (**A, B**), L-type Ca^2+^ channels current, *I*_CaL_ (**C**), the net ionic current, *I*_net_ (**D**), and *I*_Kr_ (**E**) during APs with 55% *I*_Kr_ block in heart failing model. Each simulation of block (Left) and facilitation (Right) was started from same initial values.

### Modeling suggests I_Kr_ facilitation could improve patient safety

Class III antiarrhythmic agents are powerful antiarrhythmics used to treat patients with serious ventricular tachycardias in spite of their narrow therapeutic index (*Tamargo et al. 2015*). The maximal therapeutic dose of Class III antiarrhythmics is generally limited by proarrhythmic risk of *I*_Kr_ block. At low doses with minimal proarrhythmic risk, nifekalant suppresses malignant ventricular tachyarrhythmia (*Takenaka et al. 2001*) and improves short-term and longterm survival of adult patients with ventricular fibrillation/pulseless ventricular tachycardia (*Sato et al. 2017*). In animal models, a *torsades de pointes* response was not detected with the therapeutic dose of nifekalant (*Satoh et al. 2004*), and nifekalant has a broader safety window than *I*_Kr_ blockers without facilitation (*Sugiyama 2008, Yamakawa et al. 2012*). To determine whether nifekalant’s favorable safety profile could be related to the facilitation mechanism, we conducted simulations of nifekalant’s predicted safety window with and without facilitation.

We calculated the “therapeutic” dose of nifekalant that prolongs APD_90_ by 500 ms, an established clinical standard (*Igawa et al. 2002, Drew et al. 2010*). With facilitation, the simulated therapeutic dose was 278.9 nM (68.0% *I*_Kr_ block); without facilitation, 14% less drug was required, 240.2 nM (64.1% *I*_Kr_ block) (*Figure 7*). To confirm the antiarrhythmic effect of nifekalant at those therapeutic doses, we assessed the vulnerability of cardiomyocytes to premature excitation. We measured the refractory period of APs in the normal, non-failing model by calculating the shortest timing sufficient to produce secondary voltage overshoot by a second stimuli (S2 stimuli) after regular pacing (S1-S2 stimulation protocol). The effective refractory periods with nifekalant were 216 ms or 215 ms longer than control, with or without facilitation, respectively (*Supplemental Figure 5*), suggesting that facilitation has little impact on effectiveness as a Class III antiarrhythmic. We simulated the safety window for nifekalant as the ratio of the dose that produces toxicity (EAD) to the therapeutic dose. Without facilitation, EADs occurred at 657.5 nM, 2.74× the calculated therapeutic dose (*Figure 7*). With facilitation, EADs did not occur until 894.5 nM, 3.21 × the therapeutic dose. Thus facilitation widens the simulated safety window for EADs, suggesting that with facilitation an *I*_Kr_ blocker can remain an efficacious class III antiarrhythmic agent, with an improved safety profile.

**Figure 7.**
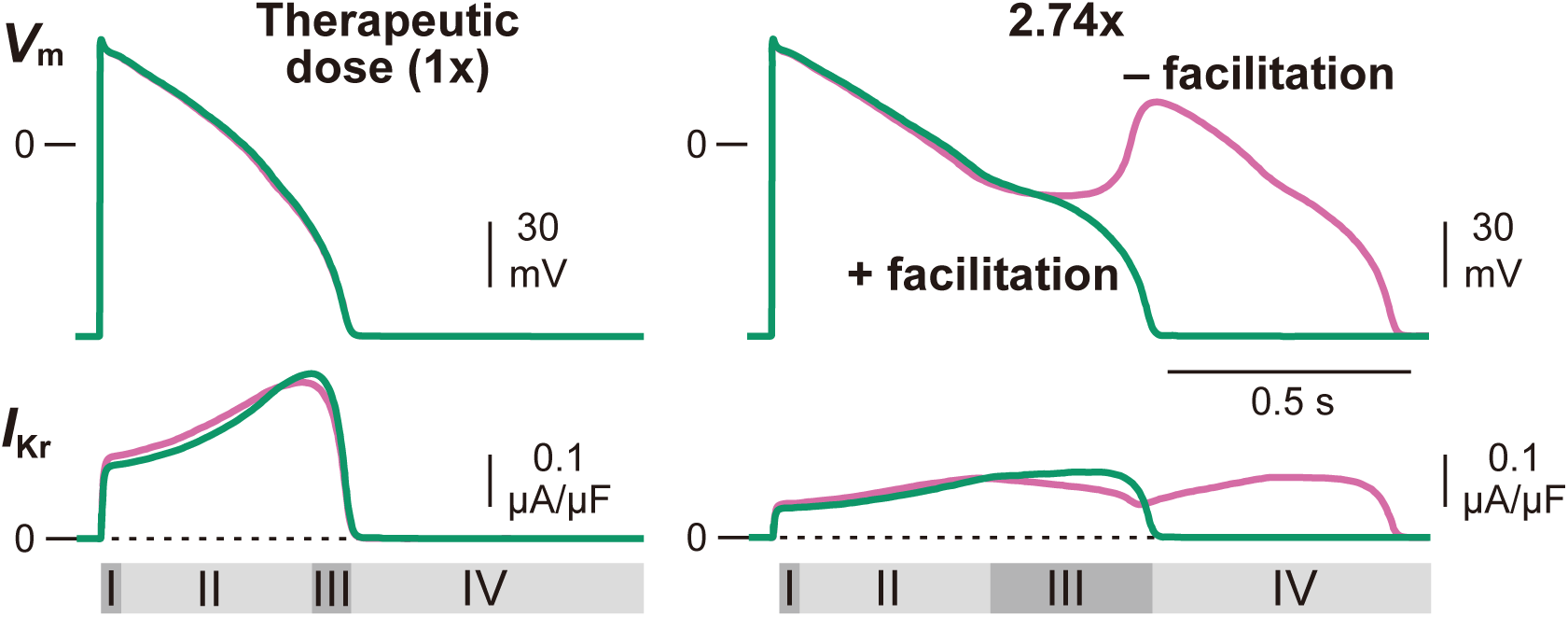
Effect of *I*_Kr_ facilitation on the safety window of nifekalant. The changes in the membrane potential (*V*_m_) (upper) and *I*_Kr_ (lower) in the presence of various concentrations of nifekalant in normal, non-failing model. The therapeutic dose is set as the concentration that prolongs APD_90_ by 500 ms and effectively suppresses the ectopic excitation (see also *Supplemental Figure 5*). 2.74× indicates 2.74 times higher drug concentration compared with its therapeutic dose (1×). Green and magenta lines indicate *I*_Kr_ block with and without facilitation, respectively. Roman numerals at the bottom indicate the phases of the AP with facilitation.

## Discussion

The objective of this study was to examine the influence of the facilitation mechanism on electrical activity in cardiomyocytes. This mathematical model represents a general view of the role that facilitation may play with *I*_Kr_ blockers. We found that *I*_Kr_ facilitation effects on simulated cardiac APs depended on the AP morphology and APD. Facilitation enhances *I*_Kr_ during prolonged APs. The repolarizing effect of facilitated *I*_Kr_ is augmented at low frequencies and as APD prolongs, thus avoiding repolarization reserve impairment, and suppressing EAD development. Therefore, hERG channel blockers with facilitation may offer a safety advantage for arrhythmia treatment.

Our conclusions are based on modeling, yet suggest a general cardioprotective mechanism. To exert cardioprotective effects *in vivo*, certain conditions are required. First, the concentration of the agent needs to reach the effective concentration for facilitation. Second, the membrane potential must induce the *I*_Kr_ facilitation effect. We have previously reported the concentration- and voltage-dependence relationships between block and facilitation for several drugs in hERG channels expressed in *Xenopus* oocytes (*Furutani et al. 2011, Yamakawa et al. 2012*). In the present study, we evaluated the concentration-dependence of nifekalant on hERG channels expressed in HEK293 cells. As in oocytes, the IC_50_/EC_50_ for block and facilitation by nifekalant were similar (92.84 ± 7.71 nM for facilitation; 144.92 ± 16.00 nM for block, see *Table 1*). However, for other class III antiarrhythmic agents, the concentration-dependences of block and facilitation are distinct. For example, the EC50 for *I*_Kr_ facilitation by amiodarone is lower than that for IC_50_ block (*Furutani et al. 2011*). These data indicate that when these agents are administrated in the treatment of arrhythmias concentrations effective for block, they certainly also reach the effective concentration for facilitation. Our experimental data showed that facilitation by nifekalant could be induced by cardiac APs (*Figure 1*). An important finding of this work is that facilitation had a negligible impact on APs under normal conditions (*Figure 3*), suggesting that facilitation would not greatly affect normal ECGs. Our simulation study suggests that facilitation may have a selective impact during severe repolarization impairment and heart failure conditions, leading to a lower risk of arrhythmias.

It is widely accepted that *I*_Kr_ plays an important role in the repolarization of the AP (*Surawicz 1989, Sanguinetti and Jurkiewicz 1990, Zeng et al. 1995, Clancy et al. 2003, Sanguinetti and Tristani-Firouzi 2006*). The decrease in *I*_Kr_ is believed to be associated with EADs, an important cause of lethal ventricular arrhythmias in LQT syndrome and heart failure. Experiments from Guo *et al* (*Guo et al. 2011*), in isolated non-failing human endocardial ventricular myocytes, showed EADs in the presence of the *I*_Kr_ blocker dofetilide (0.1 μM, corresponding to ~85% *I*_Kr_ block (*Thomsen et al. 2003*)), which induces *torsades de pointes* arrhythmias (*Thomsen et al. 2003*). In the present study using the modified ORd model, we successfully reproduced the experimental results of Guo *et al* (*Guo et al. 2011*), as did the original ORd model (*O’Hara et al. 2011*).

Reverse-frequency dependent action on APs is a property common to Class III antiarrhythmic agents (*Hafner et al. 1988, Hondeghem and Snyders 1990, Tande et al. 1990, Carmeliet 1993, Jurkiewicz and Sanguinetti 1993, Nakaya et al. 1993, Jiang et al. 1999*), and the associated proarrhythmic risk limits their clinical usefulness (*Hondeghem and Snyders 1990, Okada et al. 1996*). Reverse-frequency dependence of *I*_Kr_ block was first explained by an increase in the slowly activated delayed-rectifier K^+^ current with rapid heart rates (*Jurkiewicz and Sanguinetti 1993*). Drug mechanisms can prolong APD at higher stimulation frequencies to attenuate reverse-frequency dependence (*Hondeghem and Snyders 1990*). For example, some *I*_Kr_ blockers, such as vesnarinone, have high affinities to opened and/or inactivated hERG channels and exhibit an accumulation of inhibition at higher stimulation frequencies (use-dependent block), reducing proarrhythmic risk (*Toyama et al. 1997*). In our model, *I*_Kr_ block is not altered by stimulation frequency. Therefore, the reverse-frequency dependence may be even less than predicted. Facilitation attenuates reverse-frequency dependence by enhancing *I*_Kr_ during late phase-2 and phase-3. The effect on reverse-frequency dependence becomes more prominent at higher concentrations of drug. *Figure 4A* and *Supplemental Figure 3* shows the effects of the modest *I*_Kr_ block by 100 nM nifekalant (~40% *I*_Kr_ block and ~32% *I*_Kr_ facilitation). When the drug concentration increased, the effect of facilitation was more prominent (*Figure 4B*) because of the concentration-dependent increase in the facilitated fraction and the prolonged phase 2-3 of AP. Under the higher concentrations of blocker, EADs and alternans emerged at low frequencies that dramatically affect APD (*Figure 4B*). These EADs were effectively suppressed by *I*_Kr_ facilitation (*Figure 5*, *6*, *Supplemental Figure 4*).

Previous studies showed that *I*_CaL_ reactivation is crucial for the development of EADs (*Zeng and Rudy 1995, O’Hara et al. 2011, Tsumoto et al. 2017*). Prolongation of the time at plateau voltages caused by *I*_Kr_ block allows *I*_CaL_ reactivation (*O’Hara et al. 2011*). The EAD initiation mechanism in the present study is consistent with an *I*_CaL_ reactivation mechanism (*Figure 6*). *I*_Kr_ facilitation accelerates the repolarization just before *I*_CaL_ reactivation (*Figure 6*). Therefore, the inward-outward balance of *I*_net_ during the plateau and AP phase-3 is one determinant for the development of EADs. Such inward-outward balance of *I*_net_ results from the interaction of ionic mechanisms underlying the cardiac AP. In addition to an increase in *I*_Kr_ by facilitation, these secondary changes may contribute to preventing *I*_CaL_ reactivation and development of EADs.

Our modeling suggests that *I*_Kr_ facilitation decreases proarrhythmic risk. In considering the clinical and physiological relevance of facilitation to Class III antiarrhythmic agents, it is important to be aware of the limitations of our study. We have not yet examined the facilitation effect on native tissues. Therefore, the absolute voltage- and concentration-dependence of facilitation has not yet been determined with native *I*_Kr_ currents. Additionally, off-target effects of blocker may cause further changes, although appreciable cardiac off-target effects of nifekalant have not yet been found.

These limitations notwithstanding, this study sheds light on the potential impact of facilitation on cardiac safety, defining a novel mechanism that could suppress proarrhythmic risk and broaden the safety window of hERG blocking agents. The facilitation effect may explain the reason many clinically useful drugs, especially Class III antiarrhythmic agents, are surprisingly safe, despite hERG block and QT interval prolongation. Meanwhile, a lack of a facilitation effect may explain why dofetilide, d-sotalol, atenolol, and terfenadine have a high risk for lethal arrhythmia (*Furutani et al. 2011, Yamakawa et al. 2012*). Recent studies have described the general utility of the *in silico* integration of drug effects on multiple cardiac currents to reduce false-positive and false-negative classifications based on hERG block alone (*Mirams et al. 2011, Kramer et al. 2013*). More elaborate models incorporating detailed descriptions of ion channel modulation will improve future investigations on the clinical significance of Class III antiarrhythmic agents.

In conclusion, hERG blockers with facilitation effects may have a lower risk for inducing EADs and other triggered activities and thus be more suitable to treat arrhythmias. This finding has the potential to improve the early assessment of cardiotoxicity risk using *in vitro* ion channel assays, thereby reducing the likelihood of mistakenly discarding viable drug candidates and speeding the progression of safer drugs into clinical trials and clinical use.

## Materials and Methods

### Cell preparation and hERG channel current recording

A human embryonic kidney (HEK) 293 cell line stably-expressing hERG was kindly provided by Dr. Craig T. January (*Zhou et al. 1998*). Cells used for electrophysiological study were cultured on coverslips in 12-well plates and transferred to a small recording chamber mounted on the stage of an inverted microscope (Axiovert S100, Carl Zeiss, Oberkochen, Germany), and were superfused with HEPES-buffered Tyrode solution containing (in mM) 137 NaCl, 4 KCl, 1.8 CaCl_2_, 1 MgCl_2_, 10 glucose, and 10 HEPES (pH 7.4 with NaOH). Membrane currents were recorded in a whole-cell configuration using suction pipettes (*Hamill et al. 1981*). Leak compensation was not used. The borosilicate micropipette had a resistance of 2−4 MΩ when filled with the internal pipette solution contained (in mM) 120 KCl, 5.374 CaCl2, 1.75 MgCl_2_, 10 EGTA, 10 HEPES (pH 7.2 with KOH). Liquid junction potential with this internal solution was less than −4 mV, and the off-set was not corrected. Series resistance was typically under 5 MΩ. Series resistance compensation was used when needed to constrain voltage error to <10 mV. Whole-cell recordings were performed using an Axopatch 200B patch-clamp amplifier (Molecular Devices, Sunnyvale, CA), ITC-18 interface and PatchMaster software (HEKA Elektronik, Lambrecht, Germany). The data were stored on a computer hard disk and analyzed using PatchMaster and Igor Pro 7 (WaveMetrics, Portland, OR). In AP Clamp experiments, we used the human ventricular AP waveform simulated with our AP model as voltage-clamp commands. Most experiments were performed at a temperature of 37.0°C, which was maintained with a CL-200A temperature controller (Warner Instruments, Hamden, CT). In a few experiments, data were initially obtained with HEK293 cell and *Xenopus laevis* oocyte at room temperature as described previously (*Hosaka et al. 2007, Furutani et al. 2011*). Frogs (*Xenopus laevis*) were treated under the guidelines for laboratory animals of Osaka University Graduate School of Medicine.

Nifekalant was obtained from Nihon Schering (Osaka, Japan) and Cayman Chemical (Ann Arbor, MI).

MiRP1 (KCNE2) is a β-subunit of *I*_Kr_ (*Abbott et al. 1999*). We examined whether nifekalant caused distinct effects on currents mediated by hERG alone and by hERG and KCNE2. In these experiments, we co-expressed KCNE2 in hERG-expressing HEK293 cells by using pcDNA3 plasmid vector. However, the drugs caused virtually identical effects on the two channels currents in the dose-response (data not shown) and shift of the activation curve (*Table 1*). Therefore, we utilized HEK cells and oocytes expressing hERG channels alone for further experiments.

### Formulations of kinetic properties for hERG current

To model the macroscopic current of hERG channels expressed in HEK293 cells, we first estimated the kinetics of the channels. The voltage-dependent activation kinetics (time constant of activation) was determined from current activation experiments (*Figure 2A*). The activation time constant was measured by fitting *I*_Kr_ activation at each depolarized voltage pulse, *V*_depo_, (typically *V*_depo_ greater than −40 mV) with a single exponential function:
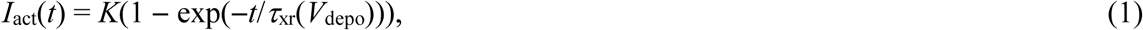

where *K* is the asymptote and *τ*_xr_(*V*_depo_) is the activation time constant. *Figure 2A* and *Supplemental Figure 1A* shows the activation time constants as a function of *V*_depo_. Furthermore, the voltage-dependent deactivation kinetics was divided into two processes (i.e., fast and slow processes) in accordance with the O’Hara-Rudy formalism (*O’Hara et al. 2011*). From current deactivation experiments (*Supplemental Figure 1B*), the deactivation time constants were measured by fitting *I*_Kr_ deactivation at each repolarized voltage pulse, *V*_repo_, (typically *V*_repo_ less than −30 mV) to a double exponential function:
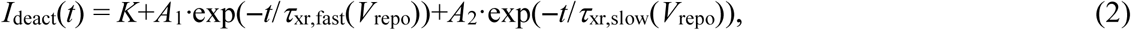

where *K* is the asymptote, and *A*_1_ and *A*_2_ are the relative components of the fast and slow processes, *τ*_xr,fast_ and *τ*_xr,slow_ are the fast and slow deactivation time constants, respectively. *Supplemental Figure 1B*, *C* show the fast and slow deactivation time constants as a function of *V*_repo_, respectively. Each time constant value in Eqs. **1** and **2** was determined by nonlinear least-squares fitting. From these experimental data, the activation and deactivation time constants were expressed as continuous functions of membrane potential, *V*_m_, as follows (*Supplemental Figure 1E*, *F*):

For *t*_xr,fast,_
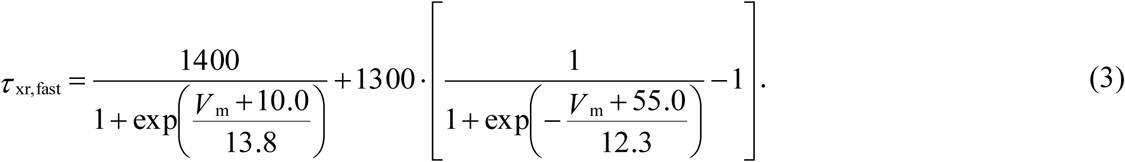

For *τ*_xr,slow,_ if *V*_m_ ≥ −80 mV
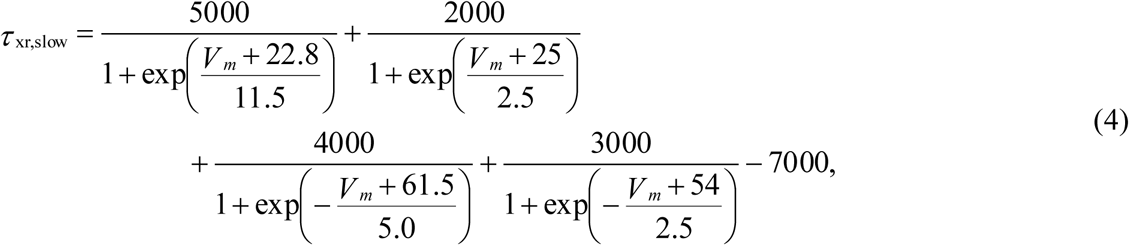

and *V*_m_ < −80 mV,
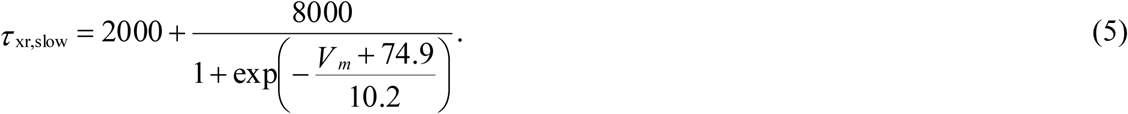

### Modeling of I_Kr_

The *I*_Kr_ current was defined as
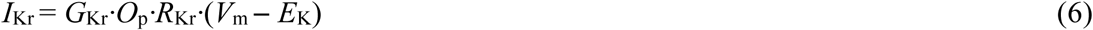

where *G*_Kr_ is *I*_Kr_ conductance (mS/cm^2^) under the drug action, *V*_m_ is the membrane potential (mV), *E*_K_ is the reversal potential, *O*_p_ is the open state variable in the activation of *I*_Kr_ channel and *R*_Kr_ is a time-independent function related to the inactivation property for *I*_Kr_, respectively. The voltage-dependence of the *O*_p_ in Eq. **6** was defined as
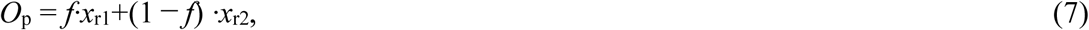

where *x*_r1_ and *x*_r2_ are voltage-dependent activation variables for unfacilitated and facilitated components in the *I*_Kr_, respectively, and *f* is a fraction of unfacilitated component in *I*_Kr_ current. The does-fraction relationship of *f* for nifekalant with an EC_50_ of 92.84 nM and a Hill coefficient (*h*_e_) of 1.50 was represented as follows:
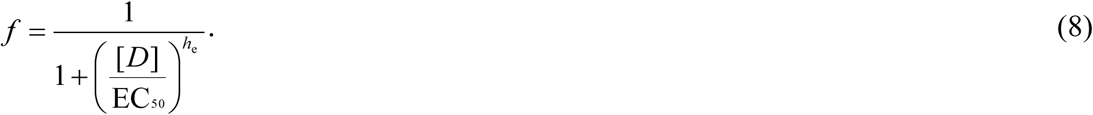

In O’Hara-Rudy dynamic (ORd) model (*O’Hara et al. 2011*), the voltage-dependent activation variable was comprised of two variables as follows:
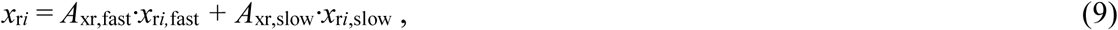

where *i* represent the voltage-dependent type, where the voltage-dependent type can be unfacilitated (*x*_r1_) or facilitated (*x*_r2_) activation variables, and *A*_xr,fast_ and *A*_xr,slow_ (≡ 1 −*A*_xr,fast_) are the fraction of channels with *I*_Kr_ activation gate undergoing fast and slow process, respectively. The fraction *A*_xr,fast_ was obtained as a following function that best reproduces voltage clamp experimental data (*Figure 2*):
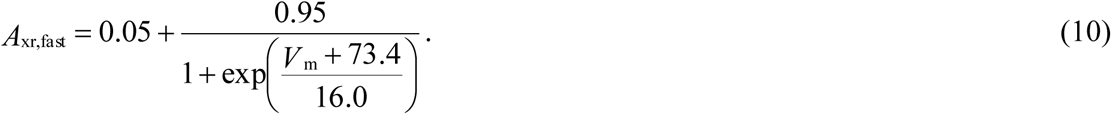

The unfacilitated (*x*_r1_) and facilitated activation (*x*_r2_) variables were calculated with the following first-order differential equation:
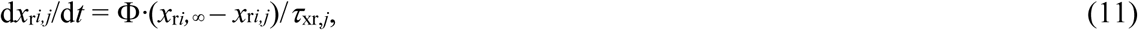

where *x*_r*i*_,_∞_, for *i* = 1, 2, is the steady-state value of *x*_r_*ij*, i.e., the voltage-dependent activation curve in the hERG channel, and *τ*_xr,*j*_ for *j* = fast, slow, is the time constant for *x*_r_*ij*, and Φ is a temperature coefficient as expressed at 3^((T-25)/10),^ where T is temperature. The voltage-dependent activation curve in the hERG channel (*x*_r*i*_,_∞_, for *i* = 1, 2) is well represented by a single Boltzmann function. Based on our experimental data (*Figure 2A* and *C*), we set the Boltzmann function’s half-activation voltage (V_1/2_) and the slope factor to –10.7 mV and of 7.4, respectively, as the voltage-dependence of the unfacilitated component (*x*_r1_,_∞_) in *I*_Kr_ activation, and shifted the voltage-dependence of facilitated components (*x*_r2,∞_) in *I*_Kr_ activation by 26.5 mV in the negative direction (*Figure 2G*), i.e.,
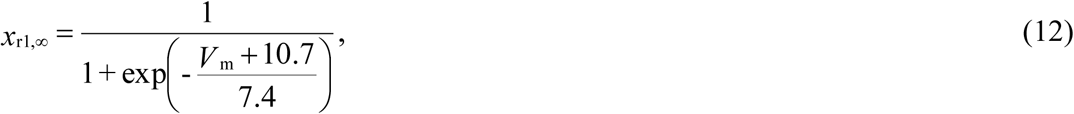

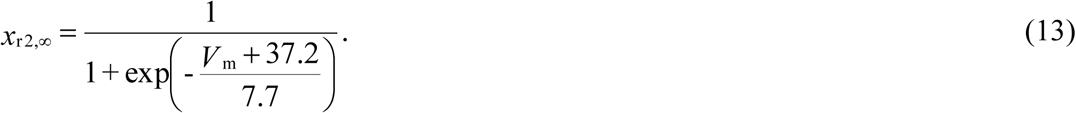

Furthermore, the *R*_Kr_ was reconstructed as a following function that best reproduces voltage clamp experimental data (*Figure 2A*):
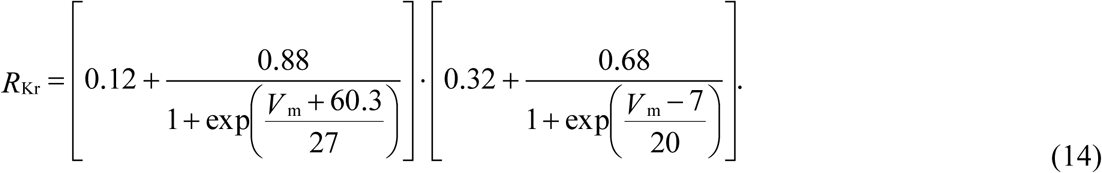

Based on our experimental measurement (*Figure 2*; *Supplemental Figure 2*), the *G*_Kr_ of when the *I*_Kr_ was blocked by nifekalant in a concentration-dependent manner with an IC50 of 144.92 nM and a Hill coefficient (*h_i_*) of 1.15 was represented as follows:
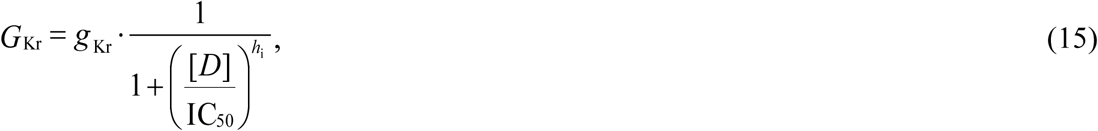

where [*D*] is the drug concentration (μM) and *g*_Kr_ is the *I*_Kr_ channel conductance absent drug. The *g*_Kr_ was set to 0.058 mS/cm^2^ to recapitulate the APD represented in the ORd model (*O’Hara et al. 2011*).

In addition, we constructed a conventional block model modified only in *I*_Kr_ conductance (block without facilitation model) for comparison with the block model with facilitation.

### Simulation protocol and computation

Voltage-clamping simulations for calculating *I*_hERG_ for each clamped voltage pulse with 4 s duration (*Figure 2*) were performed using a homemade C language program, simulating by the forward Euler method with a 0.01 ms time step. The activation variables (*x*_r*i*_, for *i* = 1, 2) was set equal to zero as an initial condition.

In AP simulations, the *I*_Kr_ model in the ORd model was replaced with our experimental based *I*_Kr_ model. The simulated AP was calculated by the fourth-order Runge-Kutta method with the double precision numbers. To minimize transient responses in each simulation, pacing stimuli of threefold diastolic threshold and the stimuli were applied repeated until the AP response observed in the ORd model reached the stationary state. To evaluate the effects of hERG channel blockade on the vulnerability of a cardiomyocyte to premature ventricular contractions, additional simulations were performed using the S1-S2 stimulation protocol: S1 stimuli until the AP converged to a steady-state were applied at the stimulating frequency of 1Hz followed by an S2 stimulus with various coupling intervals. When the arrhythmogenicity of the hERG channel blocker was examined, the stimulating frequency was set to 0.5 Hz to avoid the stimulus being applied prior to the complete repolarization of the AP. All simulations were encoded in C/C++, and run on an IBM-compatible computer with the Intel ICC compiler version 15.0.1.

### Data and materials availability

These modified ORd models (non-heart failure and heart failure models) that constructed in the present study were also implemented in an XML-based Physiological Hierarchy Markup Language (PHML), which is available at http://physiolodesigner.org/ as an open-access resource. Details on the modified ORd models as non-heart failing (and failing) endocardial myocytes with and without facilitation effect can referred from PHML models in PH database (https://phdb.unit.oist.jp/modeldb/; ID931 to 936). In addition, all AP simulations presented in this study can be reproduced by performing their PHML model simulations used a software, Flint (http://www.physiodesigner.org/simulation/flint/).

## Acknowledgments

We are grateful to Dr. M.T. Keating and Dr. M.C. Sanguinetti (University of Utah) for providing us with hERG clone and Craig T. January (University of Wisconsin) for providing us with HEK293 cell lines stably expressing hERG. We also thank Dr. Colleen Clancy and Dr. Eleonora Grandi (University of California Davis) for discussion. This study was supported by the Hiroshi and Aya Irisawa Memorial Promotion Award for Young Physiologists (to K.F. and K.T.) from the Physiological Society of Japan, Dean Award from Department of Physiology and Membrane Biology, University of California, Davis (to K.F. and J.T.S.), Grants-in-Aid for the Scientific Research on Innovative Areas 22136002 (to Y.K.), 15H01404 (to K.F.), the Scientific Research (C) 15K08231 (to K.F.), 16KT0194 (to K.T.) from the Ministry of Education, Science, Sports and Culture of Japan, the Japan Society for the Promotion of Science, and NIH grants U01HL126273 and R01HL128537 (to K.F. and J.T.S.).

## Author contributions

K.F., K.T. and Y.K. designed the experiment; K.F., K.T., I-S.C., K. H., Y. Y. conducted the experiments; K.F. and K.T. analyzed the data; and K.F., K.T., I-S.C., J.T.S., and Y.K. wrote the manuscript. All the authors revised the manuscript.

## Supplemental Figure

**Supplemental Figure 1.**
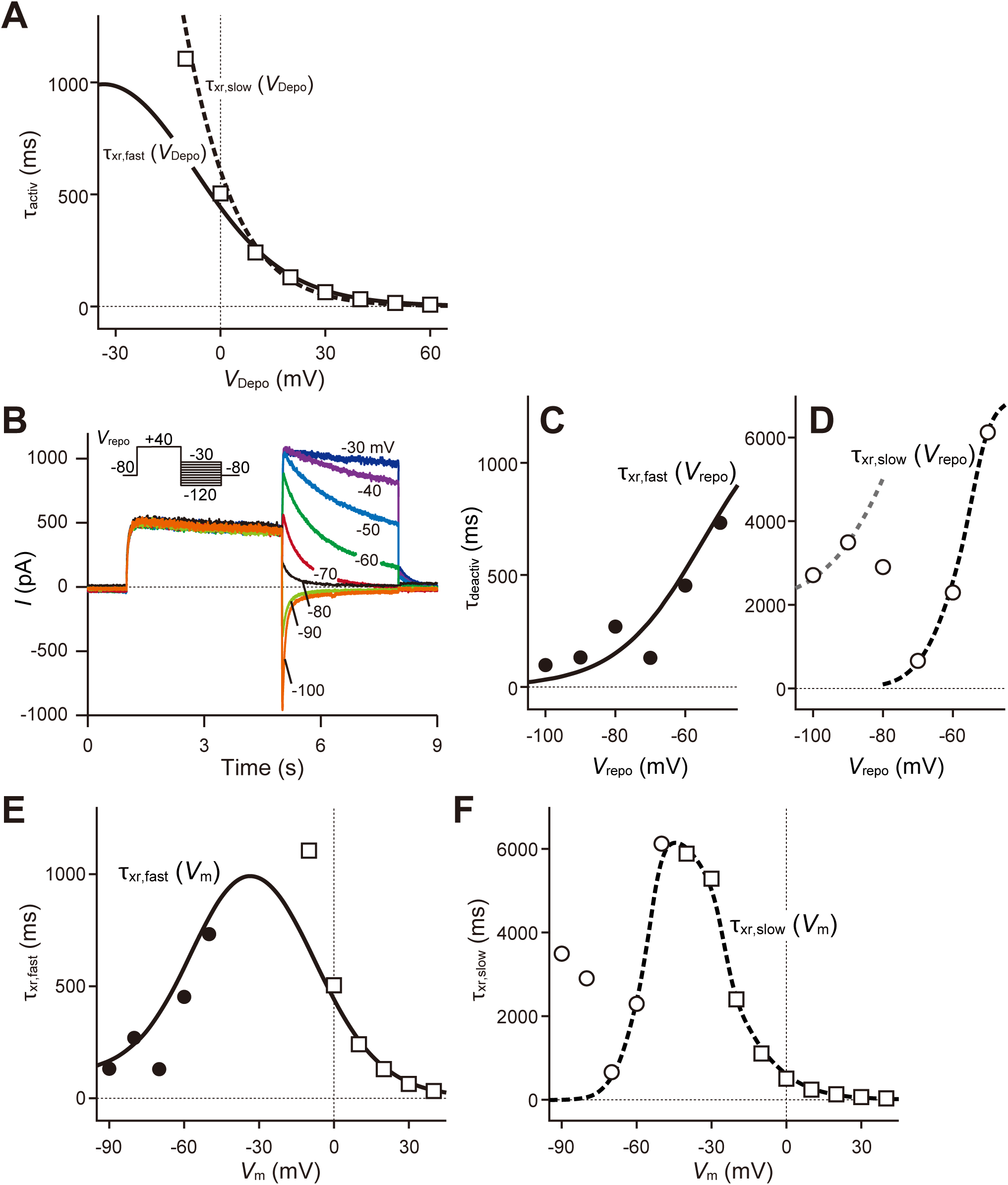
Voltage dependence of activation and fast and slow deactivation time constants. (**A**) Time constant for activation. Squares indicate activation time constant values estimated from experimental data in *Figure 2A*, and solid and dashed lines represent fast (*τ*_xr,fast_) and slow (*τ*_xr,slow_) time constants convergences, respectively. (**B-D**) Time constants for deactivation. Tail current deactivation was examined from −90 mV to −30 mV (**B**) and was fitted by a standard double-exponential equation (**c** and **d**). Points and circles represent fast (**C**) and slow (**D**) deactivation time constant values estimated from experimental data, respectively. The solid and dashed lines indicate the estimated *τ*_xr,fast_ and *τ*_xr,slow_, respectively. (**E-F**) Time constants as a function of membrane potential; fast (**E**) and slow (**F**) time constants for the activation and deactivations.

**Supplemental Figure 2.**
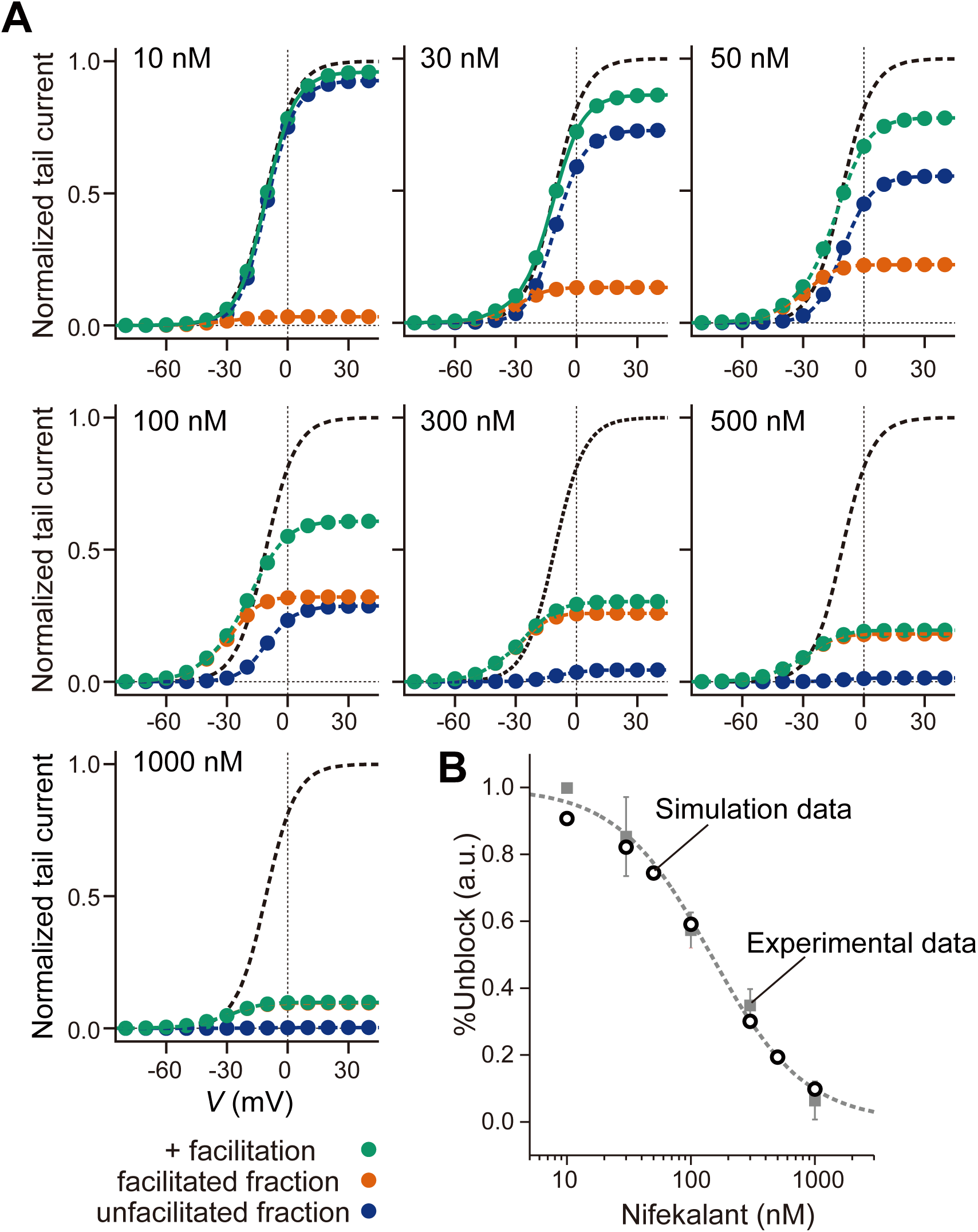
Concentration-dependent block and facilitation of nifekalant. (**A**) The relationship between the simulated tail current amplitude and membrane voltage at the indicated drug concentrations in the block with facilitation condition. Black and green lines indicate with control and block with facilitation, respectively. Orange and blue lines represent the facilitated and unfacilitated fractions of *I*_Kr_ in the block with facilitation condition, respectively. (**B**) Gray squares and gray dashed lines represent experimental concentration-dependent block and the fitted curves with Hill equation, respectively. Data are means ± SEM (*n* = 3-6). Open circles indicate the simulated concentration-dependent block. a.u. indicates arbitrary unit.

**Supplemental Figure 3.**
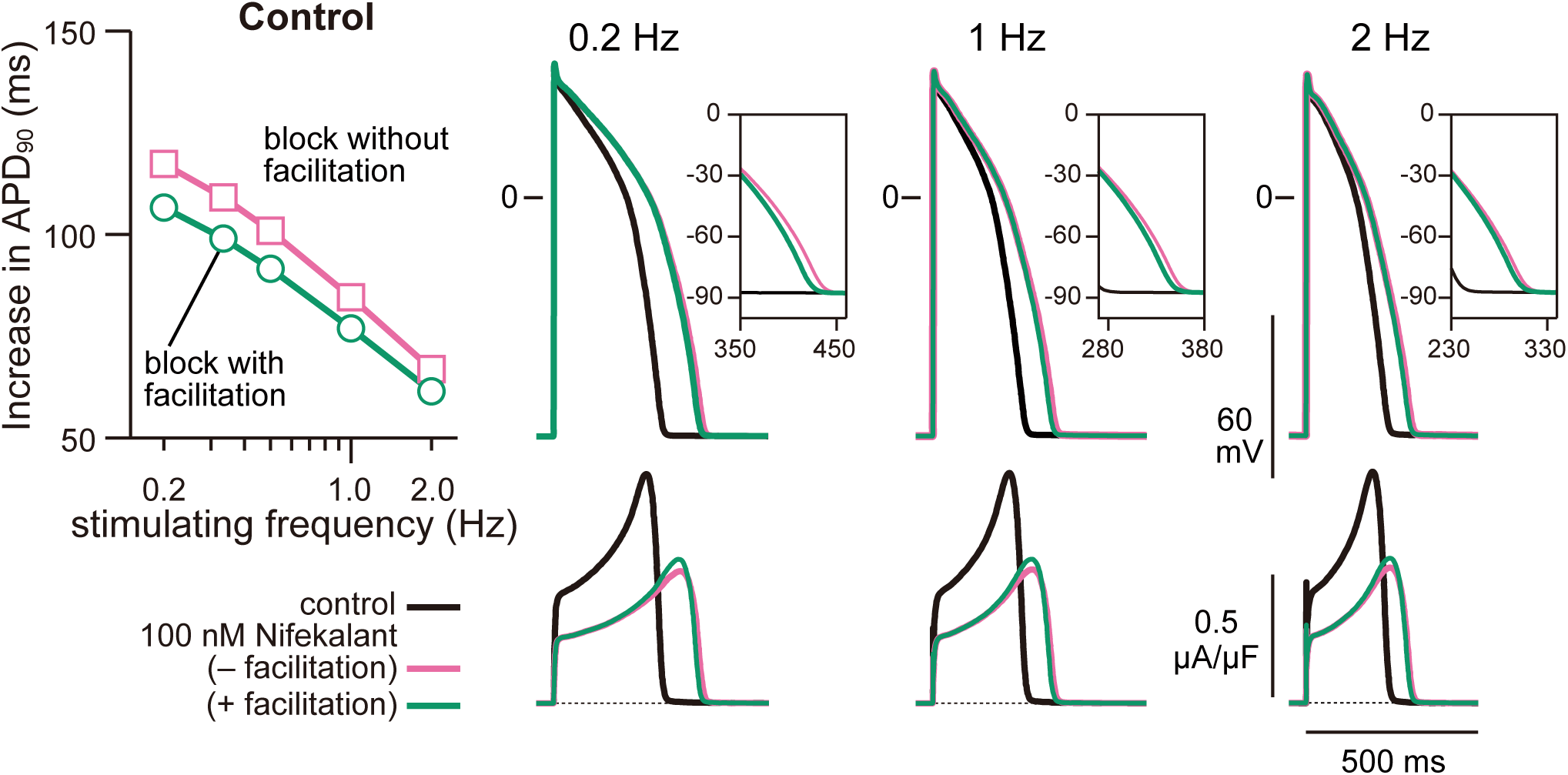
Frequency-dependent effect of nifekalant on APD in non-failing heart model. Effects of *I*_Kr_ block and facilitation on the APs in the normal, non-failing model at various simulation frequency with 100 nM nifekalant.

**Supplemental Figure 4.**
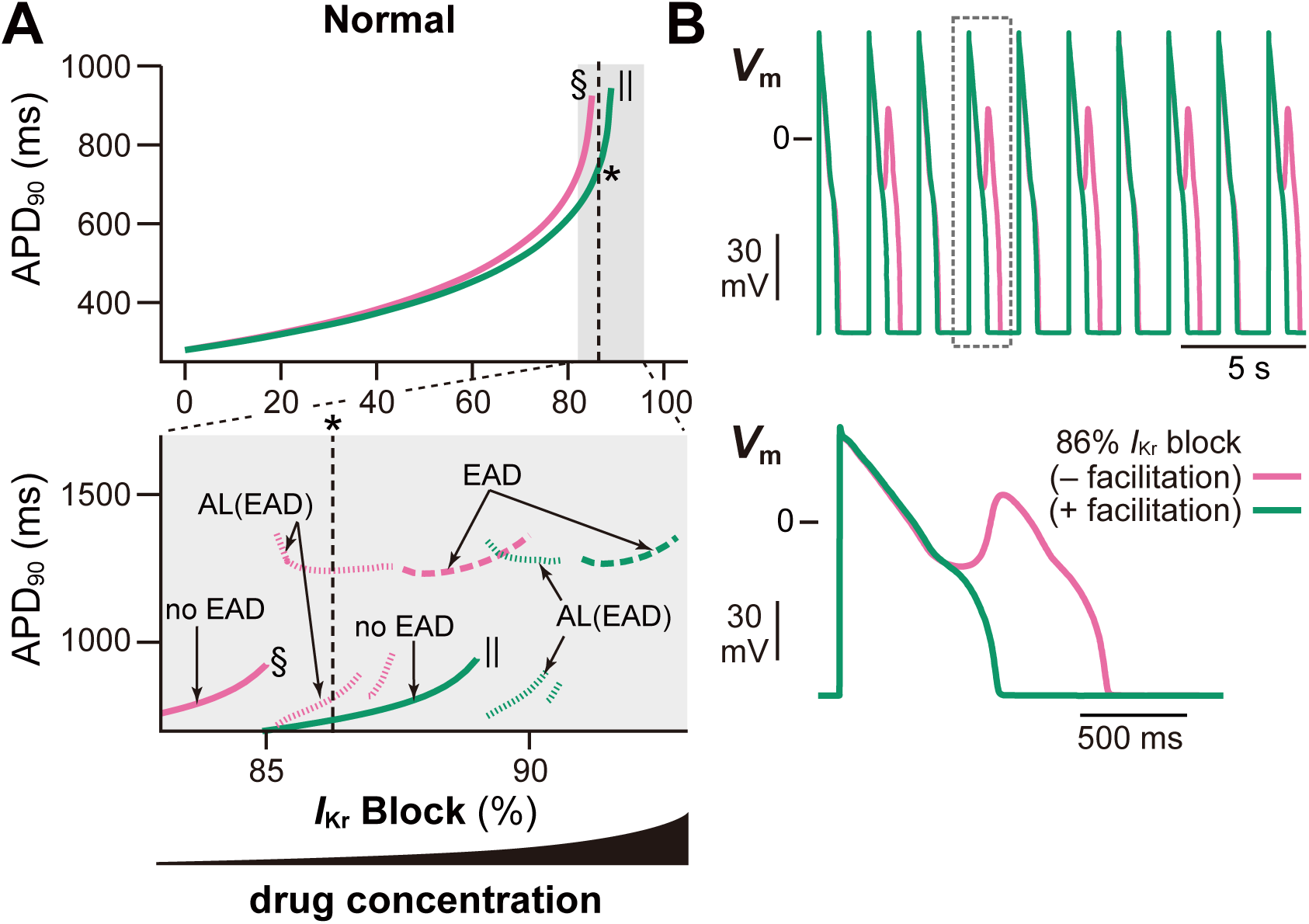
The effect of *I*_Kr_ facilitation on proarrhythmic risk in non-failing heart model. Effect of *I*_Kr_ block and facilitation on the APD and the development of EADs in normal, non-failing model (**A**). Asterisks indicate the conditions of in **B**. Symbols (daggers and double-daggers) indicate the upper limits of *I*_Kr_ block where APs ware normally terminated. (**B**) The steady-state AP trains with 86% *I*_Kr_ block in non-heart failure model with and without facilitation. Representations and symbols are the same as in *Figure 5*.

**Supplemental Figure 5.**
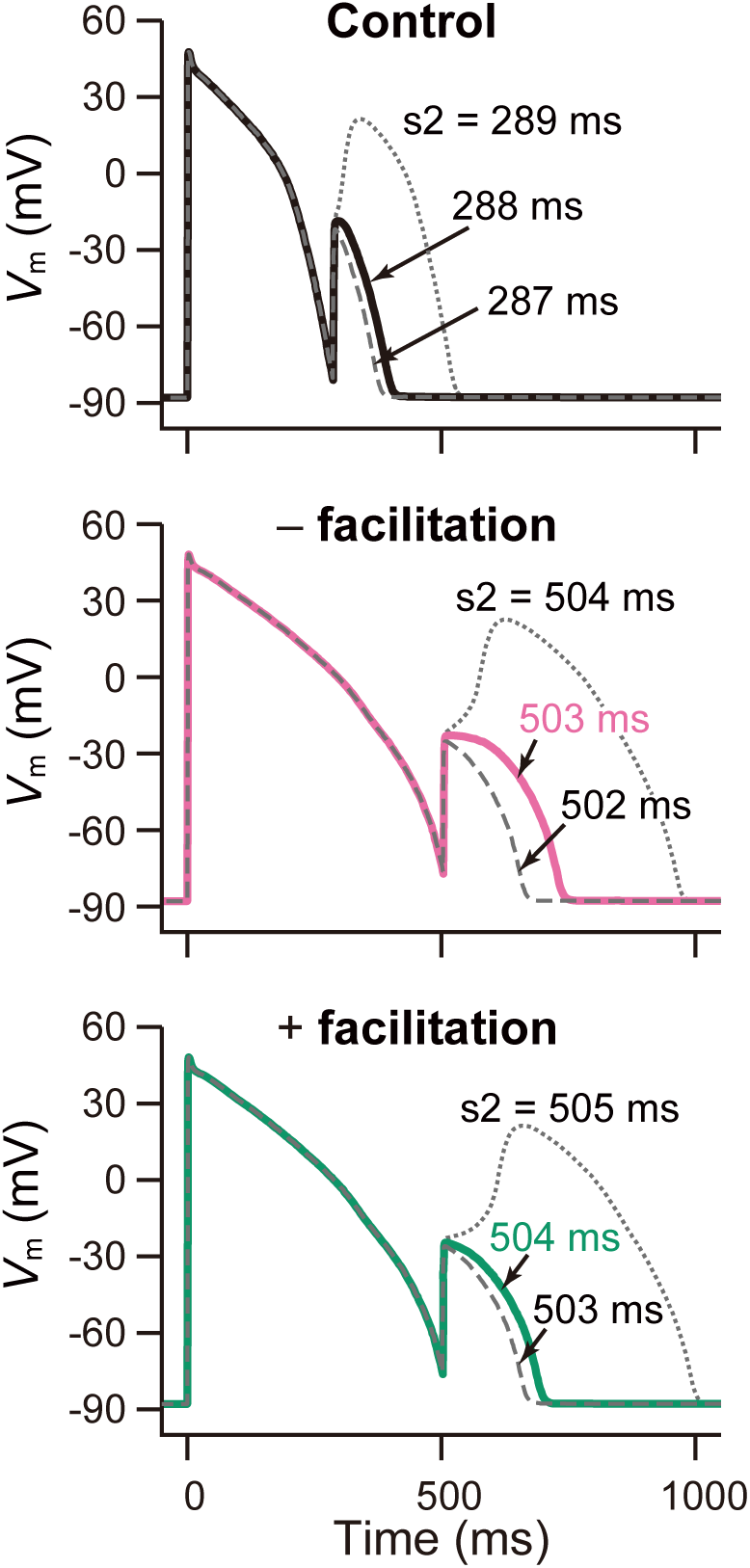
The antiarrhythmic effect of *I*_Kr_ facilitation. Simulation of block and facilitation effect on the re-excitation by the second stimulation (S2) in endocardial ventricular myocyte model with the prolonged APD_90_ by 500 ms. AP responses to the S2 stimuli applied at the time indicated.

